# Integrated Metabolic Complex Genetic Interaction Network of Chromosome 4p Loss in Basal Breast Cancer

**DOI:** 10.1101/2025.10.15.682593

**Authors:** Lynn Karam, Alain Pacis, Rohan Dandage, Michael Schwartz, Gabriel Alzial, Anastasia Gherghi, Haig Hugo Vrej Djambazian, Gerardo Zapata, Marina Schapfl, Paria Asadi, Mayerly Castrillón, Alysh Orr, Belen Hernandez, Pegah Kargar, Michelle Vandeloo, Sylvia Santosa, Andreas Villunger, Traver Hart, Jiannis Ragoussis, Geneviève Deblois, Andreas Bergdahl, Morag Park, Guillaume Bourque, Elena Kuzmin

## Abstract

Basal breast cancer subtype is enriched for triple-negative breast cancer (TNBC) and exhibits a recurrent large chromosomal deletion in chromosome 4p (chr4p). Chr4p loss is associated with poor survival, evolves early in tumorigenesis and confers on cells a proliferative state. Here, we map the integrated metabolic complex genetic interaction network of chr4p in basal breast cancer to identify targetable vulnerabilities. Differential gene expression analysis of patient derived xenografts and cancer cell models revealed that chr4p loss is associated with changes in cellular energetics and reduction/oxidation balance. Analysis of DepMap pooled genome-wide CRISPR-Cas9 screens identified complex genetic interactions specific to chr4p deletion in basal breast cancer cell models. Functional assays revealed that chr4p loss is associated with disrupted mitochondrial respiratory function and reduced glycolytic capacity, suggesting metabolic rewiring. Increased reactive oxygen species and lipid peroxidation compromised antioxidant defense mechanisms. Ultimately, this study sheds light on targeted therapies for basal breast cancer harboring large chromosomal deletions.

## Main text

Breast cancer is a heterogeneous disease characterized by diverse clinical and molecular subtypes. Therapeutic strategies have been devised for patients based on biomarkers, such as hormone (estrogen and progesterone) receptor expression or *HER2* amplification^1^. However, the basal-like breast cancer molecular subtype, constituting 10-20% of all breast cancer, often lacks these targetable biomarkers (i.e. triple negative breast cancer, TNBC) and is associated with the most aggressive behavior, the worst prognosis, leading to a large percentage of breast cancer deaths^2–4^. Basal breast cancer frequently displays recurrent large chromosomal deletions^5–7^. Structural variants occur as frequently as single gene aberrations in cancer and have been shown to arise early in tumorigenesis and confer a selective advantage^8–12^. Recent findings suggest potential therapeutic avenues for aneuploidies involving whole chromosome gains^10,13,14^. However, an understanding of the functional role and therapeutic potential of chromosome arm aneuploidies is lacking.

The most prevalent large chromosome deletions in basal breast cancer involve chromosome arms 8p, 5q and 4p^12^. Chr8p loss in breast cancer has been shown to alter fatty acid and ceramide metabolism, leading to invasiveness and tumor growth under stress conditions due to increased autophagy, thus contributing to resistance to chemotherapeutic agents^15^. It was also shown that chr5q loss in basal breast cancer leads to a loss of function of *KIBRA*, a multi-domain scaffold protein, activating oncogenic transcription factors, YAP/TAZ^16^. Our previous findings also indicate that chr4p loss occurs in ~65% of cases and correlates with poor prognosis in basal breast cancer as well as frequently observed across multiple cancer types^12^. The average deleted region on chr4p is broad encompassing a large fraction of the chromosome arm, without a minimally deleted region, and is associated with a reduced expression of ~80% of genes indicative of its functional importance. Our results further show that chr4p loss is clonal and occurs early in the course of tumor progression, indicating that it occurs in the majority of tumor cells, and confers on cells a proliferative advantage.

Genetic interactions occur when two or more mutations combine to produce a phenotype that cannot be explained by their independent effects^17^. Negative interactions occur when a combination of mutations in different genes results in a more severe growth defect than expected given the effects of individual mutations, whereas positive interactions occur when the combination of mutations confers a growth advantage. Systematic analysis of genetic interactions in model organisms has been useful for understanding the relationship between gene products at the biochemical level and how biological pathways work together to modulate cellular functions^18^. Complex genetic interactions occur when more than two genes act in concert to modulate fitness and are enriched for functionally related genes ^17,19^. Complex genetic interactions also connect functionally diverse genes, indicating that cells harboring mutations in multiple genes are perturbed for a range of cellular functions^19^. Identifying complex genetic interactions specific to chr4p loss should reveal genes that buffer its loss and shed light onto potential synthetic lethal therapeutic avenues for this poor outcome cancer.

In this study, we integrated genomic and transcriptomic analysis with DepMap pooled genome-wide CRISPR-Cas9 screens to identify genetic interactions specific to chr4p deletion in basal breast cancer models (Fig. 1). Functional assays revealed cellular changes associated with chr4p loss linking it with rewired cellular metabolism and impaired redox balance representing chr4p specific vulnerabilities.

**Fig. 1:**
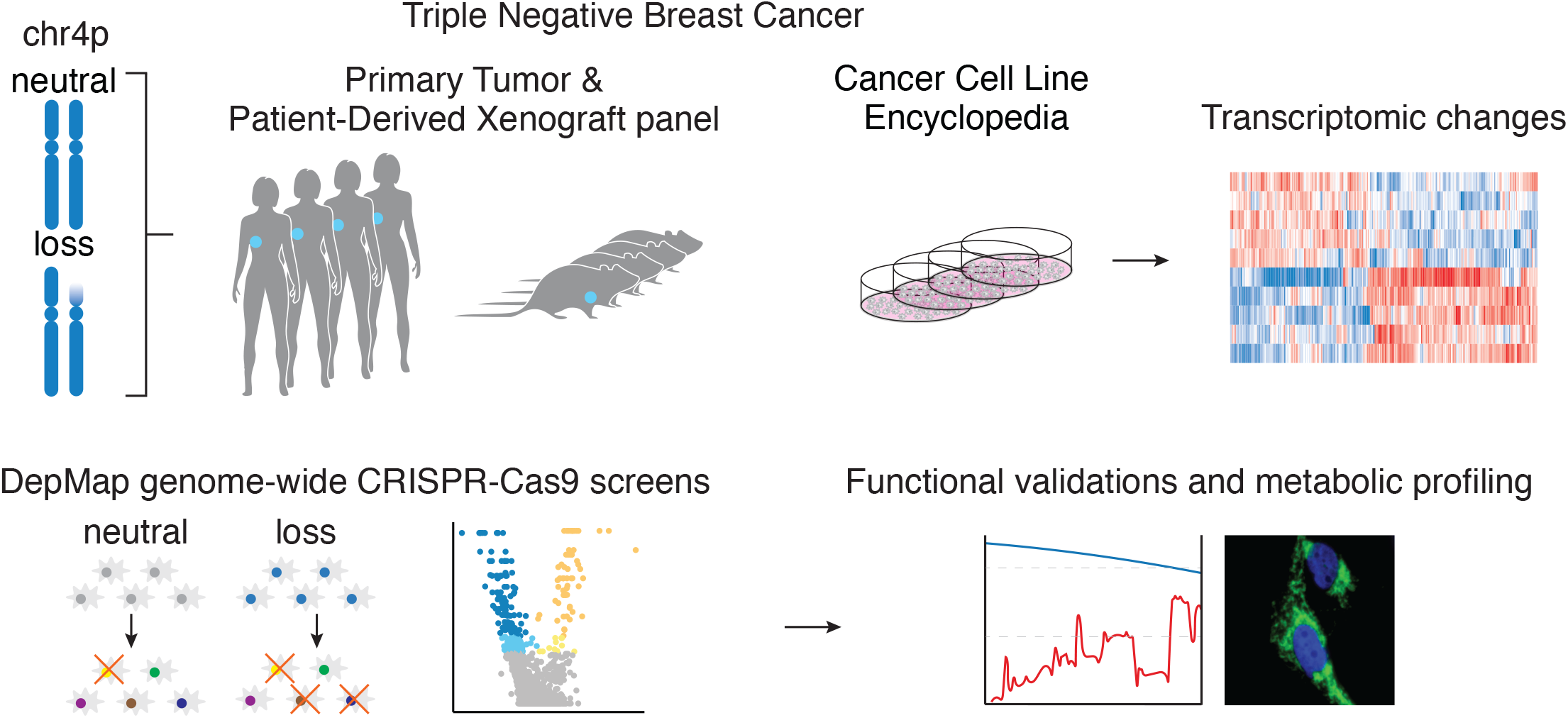
Study overview. Experimental and analytic pipeline.

## Results

### Transcriptomic changes due to chr4p loss in basal breast cancer cells

We investigated the transcriptomic changes associated with chr4p loss in basal breast cancer cells. Patient derived xenografts (PDX) established from basal breast cancer primary tumors (PT)^20^ and basal breast cancer cell lines were classified by chr4p copy number status, with chr4p loss being classified if ≥ 50% deletion of the chromosome arm was observed. Copy number log_2_ ratio was selected, which maximized the concordance between PT and PDX chr4p copy number status resulting in 12 copy neutral and 13 deletion PDX samples (Fig. 2a and Extended Data Fig. 1, Methods). Strong correlation (*r*^*2*^ = 0.93 - 0.95, *p* < 0.00001) in chr4p gene expression between PT and PDX further supports the agreement of their chr4p copy number status (Extended Data Fig. 2). Similarly, concordance between datasets from different sequencing approaches was used to classify cancer cell lines resulting in 10 copy neutral and 7 deletion samples (Fig. 2a, Methods).

**Fig. 2:**
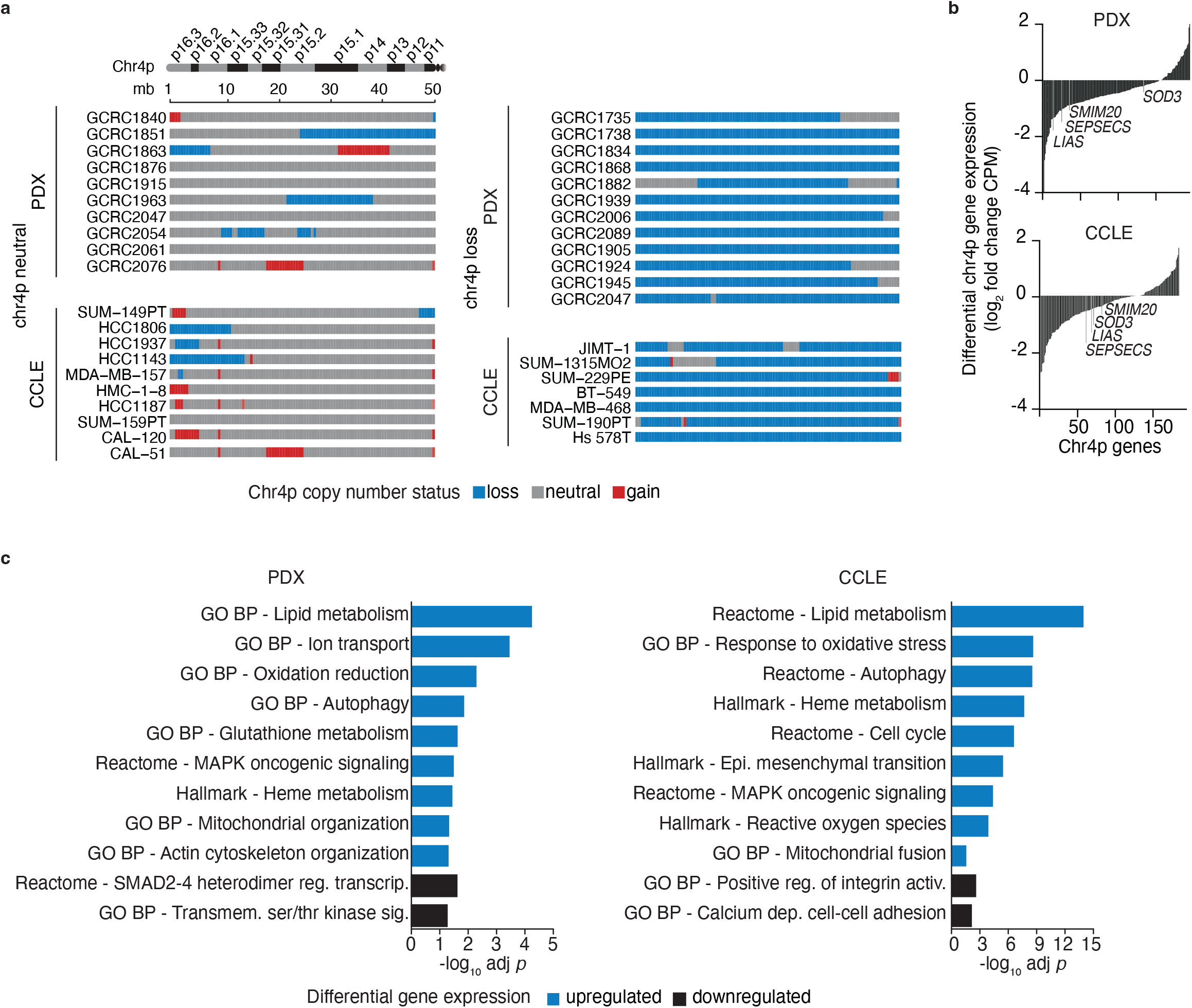
Transcriptomic changes due to chr4p loss in basal breast cancer cells. **a**, Chr4p loss and neutral basal breast cancer patient derived xenografts (PDX) and cancer cell lines using segmented mean from a previous study and the Cancer Cell Line Encyclopedia (CCLE)^64^. Log_2_ ratio ≤-0.3 shows loss in blue, ≥ 0.1 shows gain in red, and otherwise grey for neutral. Samples were classified as chr4p loss, if ≥ 50% of the chromosome arm was deleted. **b**, Log_2_ fold change of chr4p gene expression in chr4p loss compared to neutral samples across PDXs and CCLE cancer cell lines. Gene set enrichment analysis reveals differentially regulated biological pathways between chr4p loss and neutral PDXs **c**, and CCLE cancer cell lines **d**, with blue depicting upregulated and black downregulated genes.

Differential gene expression analysis of PDXs enables the capture of signal from human cancer epithelial cells unconfounded by the mouse stromal cells, which complements the analysis of CCLE cancer cell lines. Chr4p deletion was associated with a reduction in expression of ~81% (152/187) genes within the region in PDXs and ~71% (130/184) in cancer cell lines, which is consistent with a previous study^12^ (Fig. 2b and Supplementary Table 1). *SEPSECS* and *LIAS* are among the strongest downregulated genes within chr4p in PDXs and cancer cell lines. *SEPSECS* encodes an enzyme critical in the formation of the selenocysteine aminoacyl-tRNA, used to make selenoprotein which is involved in the cellular response to oxidative stress by scavenging reactive oxygen species (ROS)^21^. *LIAS* encodes an Fe-S enzyme that catalyzes the biosynthesis of lipoic acid, a potent antioxidant and is involved in copper-dependent regulated cell death^22–24^. Cancer cell lines also showed a strong downregulation of *SOD3* and *SMIM20*.

*SOD3* encodes a member of the superoxide dismutase family, which scavenges ROS from the cell and has been previously described as a potential tumor suppressor gene in breast cancer^25^. *SMIM20* encodes a mitochondrial inner membrane component of the mitochondrial translation regulation assembly intermediate of cytochrome c oxidase complex (MITRAC)^26^. The differences in the magnitude of the differential gene expression between PDXs and cancer cell lines are likely due to the effect of the 3D organization and the tumor microenvironment in the PDX, which is absent in 2D cancer cell lines. The downregulation of these genes suggests that chr4p deletion compromises mitochondrial function and the response to oxidative stress.

Chr4p deletion in basal breast cancer epithelial cells was associated with global transcriptomic changes (Fig. 2c). Basal breast cancer PDXs with chr4p loss showed an upregulation of genes with roles in lipid metabolism (*p* = 9.8 × 10^−6^), such as *HSD17B8* encoding a 17β-hydroxysteroid dehydrogenase involved in mitochondrial oxidation of fatty acids^27^. Increased lipid accumulation has been shown to be the result of dysregulation of mitochondrial oxidative metabolism combined with increased ROS production then further leading to reduced mitochondrial function^28^. Genes involved in ion transport were also upregulated (*p* = 1.6 × 10^−6^), such as *UCP2* encoding a mitochondrial uncoupling protein which separates oxidative phosphorylation from ATP synthesis, also referred to as mitochondrial protein leak, and decreases ATP production^29^. CCLE chr4p loss cell lines showed an elevated expression of genes involved in oxidative stress (*p* = 1.8 × 10^−11^), such as *RUNX1*, encoding a transcription factor, when upregulated, leads to increased oxidative stress by causing mitochondrial dysfunction, which produces excess ROS^30^. Together, these terms suggest compromised mitochondrial function and elevated oxidative stress in chr4p loss compared to copy neutral samples. Decreased expression of genes in basal breast cancer PDXs involving apoptosis regulation, such as SMAD2-4 heterodimer (*p* = 2.6 × 10^−3^), and in CCLE cell lines involving calcium dependent cell-cell adhesion (*p* = 7.6 × 10^−3^), reflects increased proliferation and a mesenchymal state associated with chr4p deletion consistent with previous findings^12^. Since genome-wide copy number profiles and average levels of aneuploidy varied similarly between chr4p loss and neutral samples, the transcriptomic changes are specific to chr4p loss and not due to another copy number change (Extended Data Fig. 1c,3).

### Systematic identification of chr4p loss specific genetic interactions in basal breast cancer

To systematically identify complex genetic interactions associated with chr4p loss in basal breast cancer, we analyzed Cancer Dependency Map (DepMap)^31^ pooled genome-wide CRISPR-Cas9 screens for 17 basal breast cancer cell lines with varying chr4p copy number status. We used a previously established computational method, MAGeCK^32^, to compare the relative abundance of gRNAs in the panel of chr4p copy neutral cell lines to the relative abundance of gRNAs targeting the corresponding genes in the panel of chr4p copy deletion cell lines (Fig. 3a). In this context, negative genetic interactions are identified as genes whose corresponding gRNAs exhibit significantly decreased abundance in chr4p deletion relative to the copy neutral background, whereas positive interactions reflect genes with increased gRNA abundance in the deletion relative to copy neutral cell lines. The log_2_ fold change represents median-summarized gRNA-level interactions. In total, we identified 217 negative and 65 positive genetic interactions associated with chr4p loss in basal breast cancer cell lines (Fig. 3b, Supplementary Table 2).

**Fig. 3:**
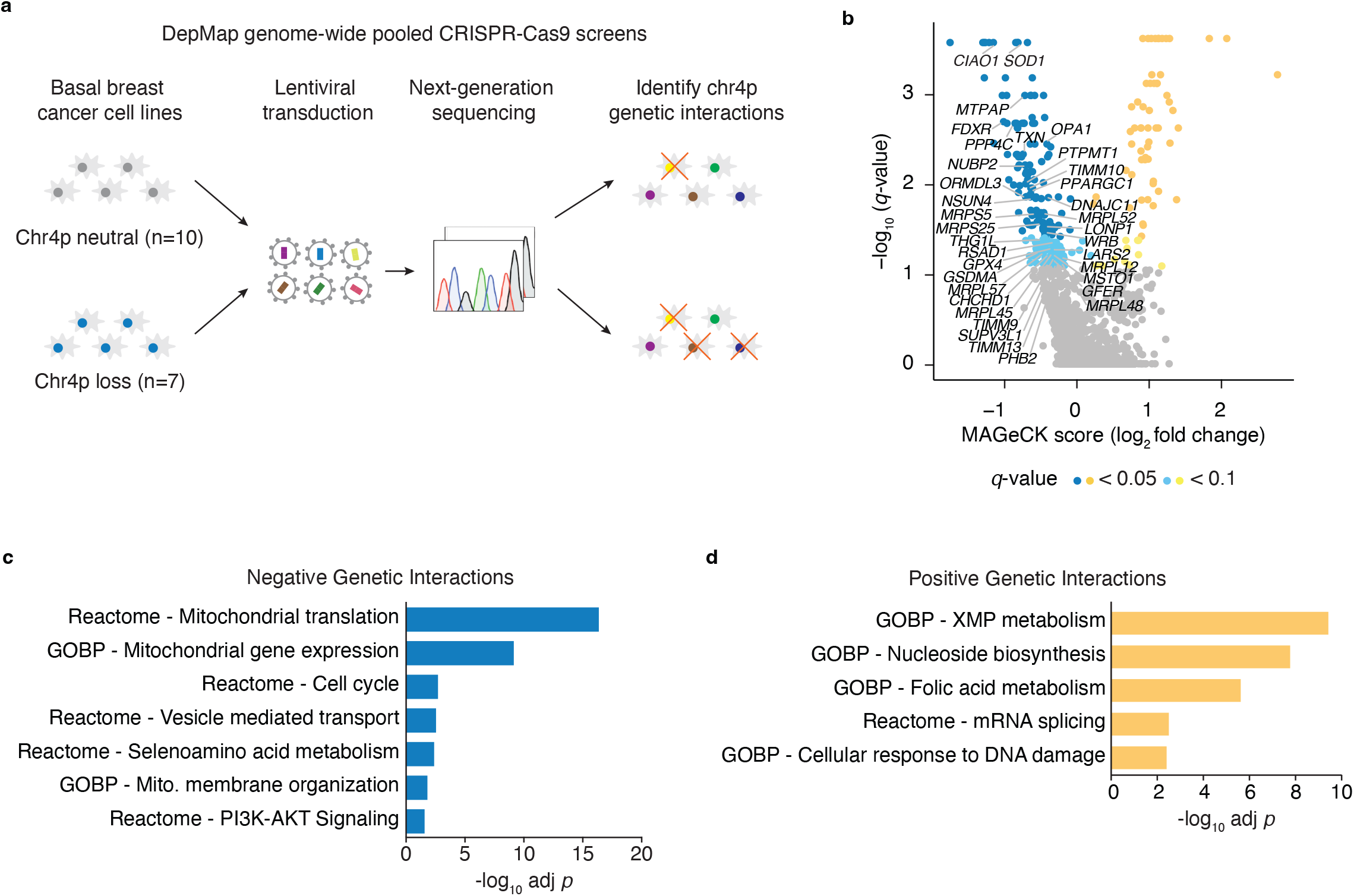
Systematic identification of chr4p loss specific genetic interactions in basal breast cancer. **a**, Schematic of DepMap pooled genome-wide CRISPR-Cas9 screening data analysis. **b**, Volcano plot depicting chr4p-specific complex genetic interactions. Negative complex genetic interactions are in blue, positive complex genetic interactions are in yellow. Different shades of these colors correspond to genetic interactions that pass different significance thresholds: *q* < 0.05 (lighter blue or lighter yellow); *q* < 0.1 (darker blue or darker yellow), grey is nonsignificant *q* > 0.1. Gene set enrichment analysis for chr4p **c**, negative **d**, and positive interactions, |MAGeCK Score log_2_ fold change| > 0, *q* < 0.1.

At a pathway level, significant negative genetic interactions were strongly enriched for genes annotated with roles in mitochondrial translation (*p* = 9.1 × 10^−21^), including mitochondrial small and large subunits, such as *MRPL52* encoding a protein involved in the translation of mitochondrial RNA^33^, mitochondrial gene expression (*p* = 8.6 × 10^−13^), such as *TRNT1* encoding the enzyme essential for modifying mitochondrial tRNAs controlling mitochondrial gene expression^34^, and mitochondrial membrane organization (*p* = 2.6 × 10^−4^), such as *DNAJC11* which is involved in mitochondrial cristae formation^35^ and *TIMM10* mitochondrial inner membrane chaperone involved in mitochondrial transmembrane transport^36^ (Fig. 3c). Negative genetic interactions were also enriched for genes involved in selenoamino acid metabolism (*p* = 4.1 × 10^−5^), such as *RPL14* regulator of selenoprotein synthesis and selenium metabolism^29^. Selenium acts as an antioxidant primarily through selenocysteine residues in ROS-detoxifying enzymes like glutathione peroxidase, thioredoxin reductase and selenoprotein P, which functions as the major transporter of selenium to tissues, and to a lesser extent through selenomethionine^37^.

Positive genetic interactions represent genes whose loss of function results in greater proliferation of chr4p loss than copy neutral cells (Fig. 3d). Chr4p positive genetic interactions involved genes annotated to such processes as xanthosine monophosphate (XMP) metabolism (*p* = 1.9 × 10^−10^). Dysregulated nucleotide metabolism has been shown to promote tumour growth^38^. Positive genetic interactions were also enriched for folic acid metabolism (*p* = 1.4 × 10^−5^). Folate deficiency was shown to lead to inefficient DNA repair and increased chromosome mis-segregation and breakage^39^. Chr4p loss aneuploidy may activate DNA damage response leading to resistance to further DNA damage, which has been previously shown for aneuploid cells with whole chromosome gains^14^. In summary, these results suggest that chr4p deleted cells depend on mitochondrial biogenesis and redox balance. Furthermore, our data illuminate the genetic determinants of how chr4p aneuploid cells are maintained in basal breast cancer.

### Chr4p negative complex genetic interactions form a metabolic functional network

To better understand the complex genetic interaction landscape of chr4p loss, we used the STRING^40^ to predict the network of functional associations between metabolic proteins encoded by chr4p negative complex genetic interactions (Fig. 4a). The resulting network involved genes encoding proteins with diverse biological functions, redox balance (*SOD1, GPX4, TXN*), iron-sulfur clusters (Fe-S) (*CIAO1, NUBP2, FDXR, RSAD1)*, mitochondrial matrix (*LONP1*), mitochondrial membrane (*PTPMT1*), mitochondria network morphology and fusion (*OPA1, MSTO1, DNAJC11, THG1L*), mitochondria biogenesis (*PPARGC1B*), mitophagy (*WRB*), translocase of the inner mitochondrial membrane (*TIMM10, TIMM9, TIMM13*), mitochondrial import (*GFER, STARD3*), ER-mitochondria contact (*ORMDL3*), mitochondrial translation (*LARS2, CHCHD1, MRPS25, MRPL45, MRPL57, MRPS5, MRPL12, MRPL48, MRPL52, NSUN4*), mitochondrial transcription (*SUPV3L1, MTPAP*), mitochondria prohibitin complex (*PHB2*) and heme chaperone (*RSAD1*). These processes are also known to crosstalk with each other, for example, *MRPL52* and *THG1L* have been implicated in protection from oxidative stress^41,42^.

**Fig. 4:**
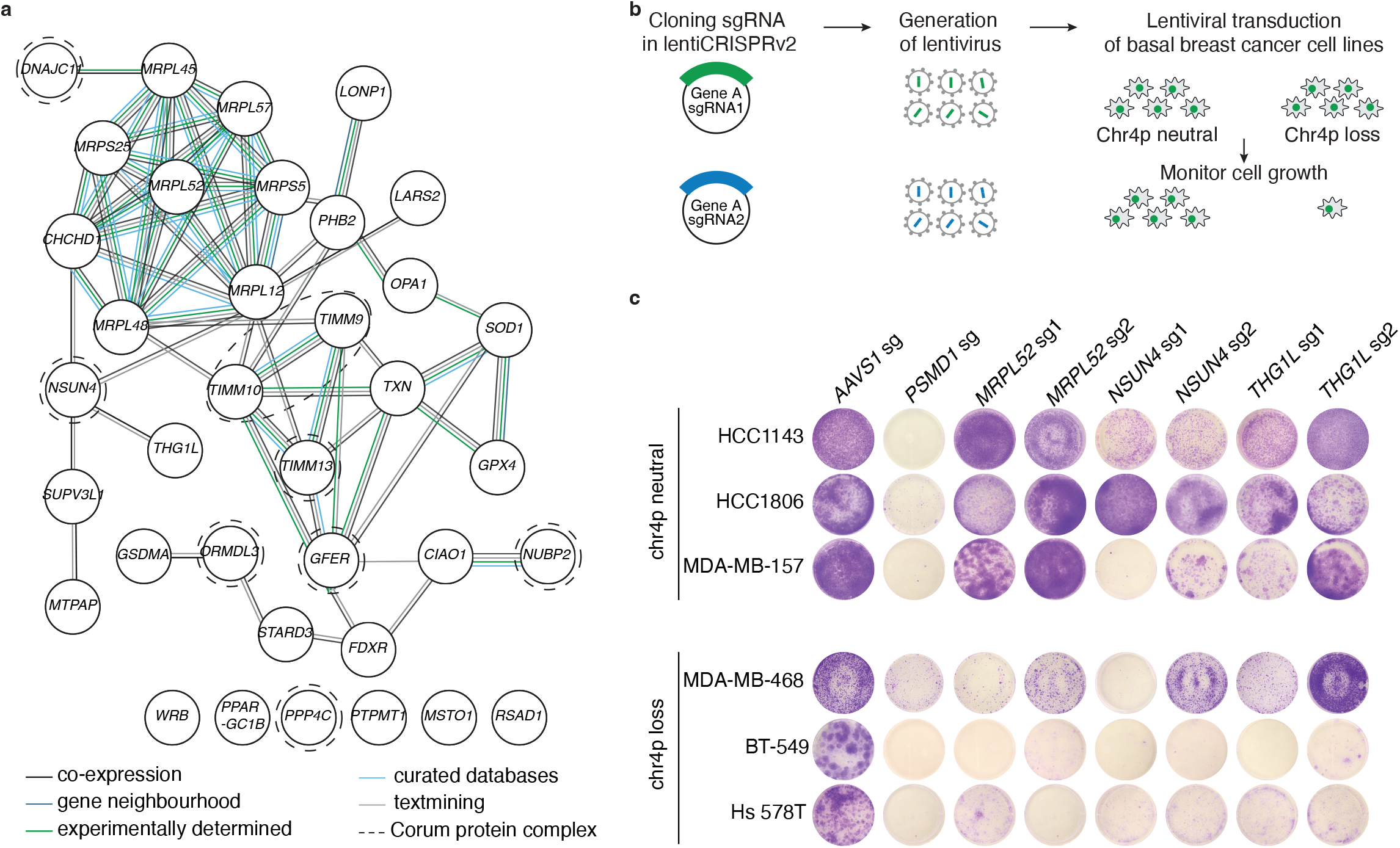
Chr4p negative genetic interactions form a metabolic functional network. **a**, STRING ^40^ network showing chr4p negative genetic interactions (MAGeCK Score log_2_ fold change < 0, *q* < 0.1) involved in mitochondria function. Nodes represent proteins produced by a single protein coding gene locus and edges represent protein-protein interactions, co-expression and gene neighbourhood. Experimentally determined and database curated interactions are represented in light and medium blue, predicted interactions are in dark blue, gene neighborhoods and other interactions are in black and co-expressed and text-mined interactions are in grey. Dashed outline highlights a CORUM^43^ protein complex capturing ≥50% members. **b**, Schematic of the genetic interaction validation strategy. **c**, Crystal violet stained wells of cells targeted by CRISPR-Cas9 sgRNA in chr4p neutral and loss basal breast cancer cell lines. sgRNA targeting *AAVS1* and *PSMD1* represent a negative and positive control, respectively.

A further analysis of the STRING network of negative genetic interactions with the CORUM protein complex standard^43^ revealed members of the small translocase of the inner membrane (TIM) chaperone complex (*TIMM9, TIMM10, TIMM10B*) and TIMM8A-TIMM13 complex (*TIMM13*) mediating the import and insertion of multi-pass transmembrane proteins into the mitochondrial inner membrane^44^, *DNAJC11* of the mitochondrial intermembrane bridging (MIB) complex involved in the formation and maintenance of mitochondrial cristae affecting the assembly of respiratory complexes^35^ and *NSUN4* of the MTERF4-NSUN4 complex, which is involved in mitochondrial ribosomal biogenesis^45^. We also identified negative genetic interactions which encoded proteins with roles in redox balance, such as *ORMDL3* of the ORMDL3-SPTLC1 complex involved in sphingolipids biosynthesis and homeostasis, which when perturbed can lead to an increase in ROS levels disrupting mitochondrial redox balance^46^ and *NUBP2* of the NUBP1-NUBP2 complex, which mediates *de novo* assembly of Fe-S cluster involved in redox reactions and ferroptosis^47^. We tested a random set of negative complex genetic interactions belonging to the metabolic network by independent validation assays and confirmed all three by examining the colony formation of a subset of three chr4p copy neutral and three chr4p deletion basal breast cancer cell lines expressing gRNAs against *MRPL52, NSUN4* and *THG1L* (Fig. 4b,c and Supplementary Table 2). Overall, our findings indicate that chr4p loss cells are perturbed for mitochondrial function and ROS detoxification resulting in their dependencies on these pathways.

### Chr4p loss is associated with metabolic rewiring

To investigate negative genetic interactions associated with chr4p loss related to metabolism and mitochondrial function, we employed a high resolution respirometry assay and measured the changes in oxygen flux of our panel of cell lines exposed sequentially to exogenous substrates (Fig. 5a). Chr4p loss cells exhibited lower routine respiration compared to copy neutral cells (Fig. 5a). Chr4p loss cells also showed higher leak respiration upon the addition of malate, glutamate and pyruvate, which are mitochondrial respiratory complex I activators capturing the portion of electron transport chain activity, which is not coupled to ATP synthesis through the process of oxidative phosphorylation (Fig. 5a). Consistent with this finding, protein abundance of mitochondrial respiratory complexes I-V was reduced in chr4p loss compared to copy neutral basal breast cancer cells (Fig. 5b). Mitochondrial complex I has been shown to be fundamental for the assembly of mitochondrial respirasomes, which are large, supramolecular assemblies of respiratory complexes of the human respiratory chain^48^. Thus, the decrease of complex I protein abundance affects the stoichiometry of other respirasomes further impairing mitochondrial function. Chr4p loss was also associated with elongated mitochondria (Fig. 5c), as evident from the immunofluorescent staining of *TOMM20*, encoding a key component of the translocase complex^49^, and upregulation of genes involved in mitochondrial fusion (*p* = 3.5 × 10^−3^), which has been shown to reduce the susceptibility of cells to apoptotic death^50^. Some metabolic heterogeneity in this subset of the tested cell lines was observed, which could be due to the individual variation among their genetic backgrounds. These functional assays combined with differential gene expression analysis and negative genetic interactions involving mitochondrial biogenesis and organization genes suggest that chr4p loss is associated with perturbed mitochondrial respiration.

**Fig. 5:**
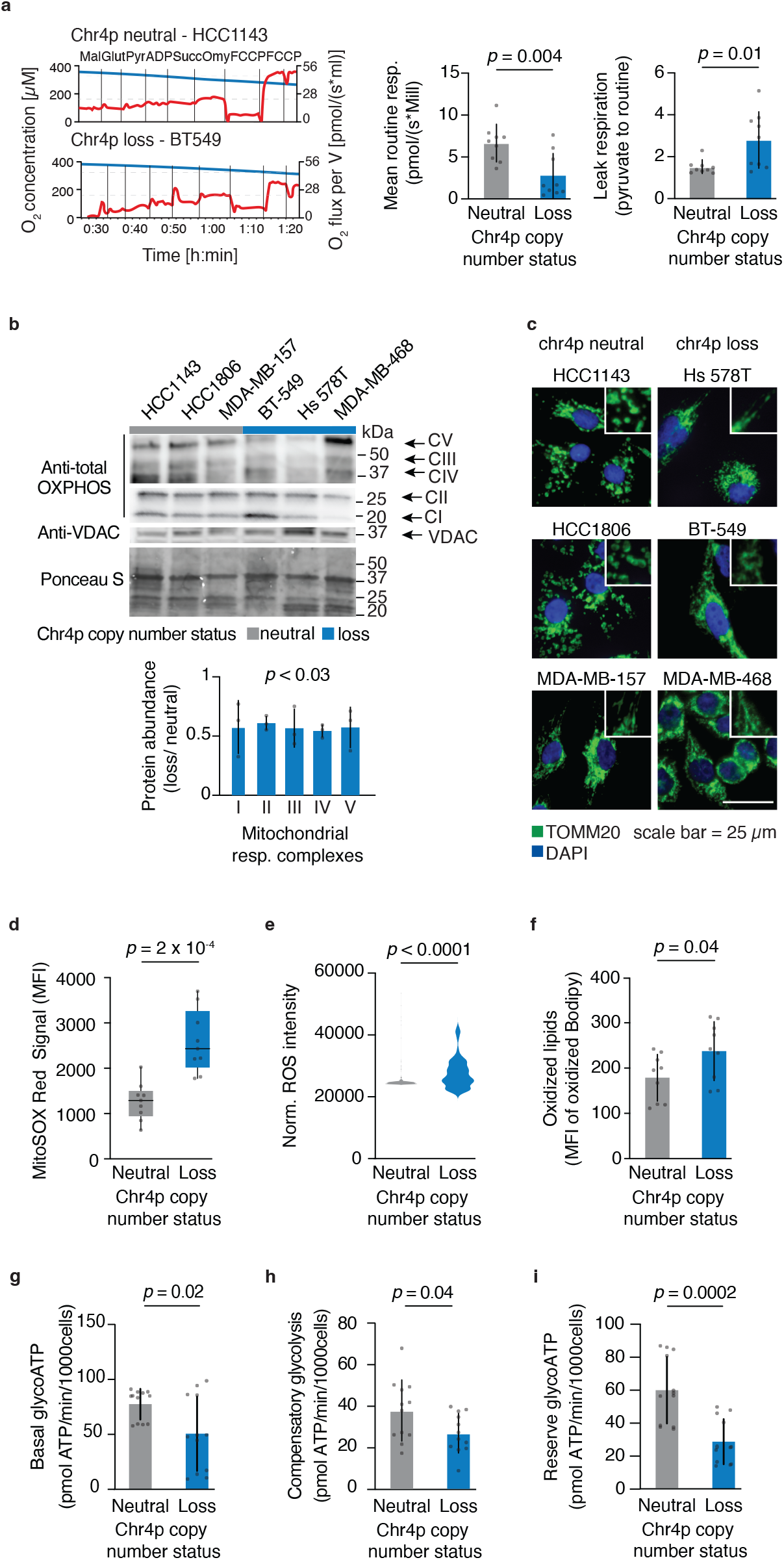
Chr4p loss is associated with metabolic rewiring. **a**, Representative plots generated by Oroboros Oxygraph 2k depicting oxygen consumption rate (red) and oxygen concentration (blue) over time for chr4p loss (BT-549) and neutral (HCC1143) cell lines. Routine respiration varies for chr4p loss and neutral cells. Mean is shown; error bars denote SD. Significance was assessed by an unpaired t-test. n=3. **b**, Western blot using whole-cell lysate shows variation in mitochondrial respiratory complexes abundance between chr4p loss and neutral cells. Mean is shown; error bars denote SD. Significance was assessed by an unpaired t-test. n=3. Exposure time for VDAC was 5 seconds, for complex I and II was 2 seconds, and for complex III-V was 60 seconds. Ponceau S stain was used for loading control. **c**, Elongated mitochondrial network in chr4p loss compared to neutral cell lines was monitored by TOMM20 immunofluorescent staining. n=3. **d**, Superoxide levels were increased in chr4p loss cells as monitored by MitoSox Red signal. Significance was assessed Wilcoxon rank sum test. n=3 **e**, Cellular reactive oxygen species were elevated in chr4p cells as monitored by CM-H_2_DCFDA probe. Significance was assessed by Wilcoxon rank sum test. n=3. **f**, Lipid peroxides were increased in chr4p loss cells as indicated by BODIPY C11 591/581 sensor. Significance was assessed by an unpaired t-test. n=3 **g**, Basal glycolytic ATP, **h**, Compensatory glycolysis, **i**, Glycolytic reserve was reduced in chr4p loss cells. Significance was assessed by an unpaired t-test. n=3. Chr4p loss (blue), neutral (grey).

To investigate the consequence of perturbed mitochondrial respiration in chr4p deletion cells, we measured the levels of diverse ROS species in our panel of basal breast cancer cell lines. We used MitoSOX red, a fluorogenic indicator, which is specifically targeted to mitochondria in live cells, to measure mitochondrial superoxides (Methods). We detected higher levels of mitochondrial superoxides in chr4p loss than copy neutral cells (Fig. 5d). An increase in mitochondrial superoxide levels was previously shown to impair mitochondrial function by disrupting the electron transport chain, damage mitochondrial DNA and proteins, and ultimately lead to a decrease in ATP production^51^. Mitochondrial superoxides are degraded through mitochondrial superoxide dismutase namely *SOD2* which transforms superoxides to hydrogen peroxide subsequently detoxified by enzymes, such as catalase in the peroxisome and glutathione peroxidase in the mitochondria^51^. *SOD2* gene expression is significantly reduced in PDXs with chr4p loss compared to copy neutral which could contribute to a reduced capacity of chr4p loss cells to transform superoxides and detoxify the mitochondria (Supplementary Table 1). We then sought to find out whether mitochondrial superoxides leak into the cytoplasm by measuring overall cellular ROS levels. We used CM-H_2_DCFDA, a chloromethyl derivative of H_2_DCFDA, which is a cell permeant indicator fluorescing upon oxidation (Methods). Cells harboring chr4p deletion exhibited increased CM-H_2_DCFDA fluorescence suggesting increased cellular ROS levels compared to copy neutral cells (Fig. 5e). Finally, we used BODIPY 581/591 C11, which is a lipophilic sensor that localizes to membranes in live cells enabling analysis of lipids oxidized by free radicals from ROS (Methods). We observed a greater shift of BODIPY fluorescence suggesting a greater level of lipid peroxidation in chr4p loss compared to copy neutral cells (Fig. 5f). Oxidative degradation of cellular lipids by ROS causes damage to cell membranes leading to aberrant signal transduction pathways and has been implicated in playing a role in cancer^51^. Lipid peroxidation in PDXs is suggested by the upregulation of genes involved in glutathione metabolism (*p* = 3.5 × 10^−4^), such as *GSTM2* encoding a glutathione S-transferase and protects against lipid peroxidation^52^ (Fig. 2c). Together these findings are consistent with misregulated expression of genes involved in ROS scavenging, glutathione metabolism and negative genetic interactions with genes involved in selenoamino acid metabolism and redox balance suggesting that chr4p loss is associated with oxidative stress in basal breast cancer.

Since chr4p loss is associated with reduced mitochondrial respiration, we performed Seahorse Cell Mito Stress assay which enables to separately investigate mitochondrial and nonmitochondrial respiration representing oxygen consumption by cellular processes outside of mitochondria^53^. Consistent with the high-resolution respirometry analysis, chr4p loss was associated with a decreased oxygen consumption rate (OCR), mitochondrial ATP production and coupling efficiency, a measure of how efficiently the energy released from electron transfer is utilized to generate a proton gradient across the inner mitochondrial membrane (Extended Data Fig. 4). Some differences in the assays are likely due to the high-resolution respirometry analysis being performed in maximal respiration conditions. We observed a reduction in glycolytic ATP production and compensatory glycolysis indicated by a lower increase in glycolytic activity in response to impaired mitochondrial ATP production by subjecting cells to modulators of respiration in chr4p loss compared to copy neutral cells (Fig. 5g-i). Once the mitochondria was impaired, chr4p loss cells did not show an increase in glycolytic ATP levels compared to basal levels (Fig. 5g,h), which is also illustrated by a lower glycolytic reserve in chr4p loss compared to neutral cells (Fig. 5i). The downregulation of a chr4p gene *PGM2*, which encodes phosphoglucomutase and catalyzes the reversible conversion of glucose 1-phosphate to glucose 6-phosphate, which is a key step in glycolysis contributes to the low glycolytic capacity associated with chr4p loss^54^ (Fig. 2b and Supplementary Table 1). These findings suggest that basal breast cancer cells harboring chr4p deletion are associated with metabolic rewiring.

## Discussion

Here, we mapped the complex genetic interaction network of chr4p loss in basal breast cancer and identified the resulting vulnerabilities. An integrated analysis of transcriptomic changes associated with chr4p deletion and complex genetic interactions identified from DepMap pooled genome-wide CRISPR-Cas9 screens revealed misregulation of metabolic and redox homeostasis pathways. Through multiple approaches, we showed that chr4p loss is associated with impaired mitochondrial and cellular ROS scavenging as well as perturbation in mitochondrial respiration and glycolytic metabolism, revealing a state of metabolic rewiring associated with this chromosome arm aneuploidy. More generally, this analysis enhances our understanding of genetic dependencies of chromosomal arm losses opening avenues for targeted therapies for TNBC.

We conducted transcriptomic and genetic screen analyses and identified genetic dependencies and interactions specific to chr4p loss in basal breast cancer, many of which were involved in mitochondrial respiration and redox balance. Perturbation of genes involved in these processes, specifically components of the mitochondrial electron transport chain as well as major antioxidant mechanisms, residing within chr4p resulting from the hemizygous deletion of this chromosome arm reflects their dosage sensitivity underlying chr4p synthetic lethal interactions. Pathways involved in oxidative stress, such as glutathione metabolism and response to oxidative stress were also upregulated in chr4p loss compared to copy neutral samples as evident from the global transcriptomic profiles. These observations likely reflect a compensatory mechanism in response to increased oxidative stress and are consistent with previous findings in aneuploid disomic yeast, with increased chromosome numbers^55^. This finding also suggests that the oxidative stress response is conserved between yeast and human cells and is common to specific whole chromosome gains and chromosome arm losses. Recent CRISPR screens of breast cancer-associated aneuploid human mammary epithelial cells with chromosome gains revealed dependencies related to mitochondrial oxidative phosphorylation^56^. In addition to coding genes, long noncoding RNA and microRNA are located on chr4p and have targets elsewhere in the genome, thus their loss may contribute to transcriptomic changes observed from chr4p deletion underlying some chr4p genetic interactions which could be explored in the future. Together, these finding suggest of converging mechanisms that buffer whole chromosome gain and chromosome arm losses.

Cancer cells have been known to maintain their high ROS levels below the threshold where apoptosis, ferroptosis or senescence are triggered^51^. Increased ROS levels causing DNA damage ultimately provide a selective advantage to aneuploid cells through p53 activation maintaining genomic stability in the presence of aneuploidy^57^. Oxidative stress may be beneficial to cancer cells as high levels of ROS can promote a proliferative state by inhibiting the activity of the tumor suppressive protein PTEN and consequently activating the oncogenic MAPK signaling pathway^51^. Furthermore, mitochondrial ROS has been shown to increase micronuclei collapse through oxidizing and activating the autophagic protein p62 and interfering with CHMP7, which is a nuclear membrane repair protein ^58,59^. By removing micronuclei, aneuploid cells with high ROS suppress innate immune detection and limit cytosolic DNA stress, which might otherwise lead to senescence or cell death. It has been shown that chr4p loss is associated with a proliferative transcriptional program^12^, which is consistent with their elongated mitochondrial network, which has been linked to reduced apoptosis susceptibility by reducing apoptotic priming thus increasing the threshold that is required to induce apoptosis^50^. Altered priming is further supported by chr4p loss PDXs showing an upregulation of genes involved in heme metabolism (*p* = 3.8 × 10^−3^) (Fig. 2c, Supplementary Table 1). Heme is important to maintain electron transport chain activity and heme biosynthesis upregulation has been implicated with metabolic rewiring associated with reduced mitochondrial respiration, decreased apoptotic priming and resistance to a proapoptotic small molecule, venetoclax, in multiple myeloma^60^, a compound which is also being investigated for triple negative breast cancer.

Compounds targeting mitochondria function have emerged as a prominent therapeutic approach for multiple cancer types. For example, mitochondrial complex I inhibitors, ME-344 and metformin, are being tested in breast cancer phase 1 and phase 2 trials, respectively ^61,62^. Since mitochondrial respiration and glycolysis are reduced in chr4p deletion cells, they rely on alternative pathways to meet their energetic demands. Consistent with this, chr4p loss PDXs showed an upregulated expression of genes involved in lipid metabolism (*p* = 9.8 × 10^−6^), such as *HSD17B8*, which has been shown to reroute 3R-hydroxyacyl-CoA esters into the mitochondrial β-oxidation to avoid accumulation of polyunsaturated fatty acids in the mitochondria^27^ and genes involved in regulation of autophagy (*p* = 1.7 × 10^−4^), such as *ACER2*, encoding a Golgi ceramidase, which has been shown to induce autophagy through the generation of sphingosine and ROS ^63^ (Fig. 2c, Supplementary Table 1).

In summary, we present an integrated, unbiased and genome-wide approach for uncovering genetic vulnerabilities related to chr4p deletion in basal breast cancer providing a resource for studying metabolic rewiring and cancer disease phenotypes linked to chromosome arm aneuploidies. Our metabolic profiling led us to identify a global decrease in mitochondrial respiration, including impaired oxygen consumption rate, mitochondrial ATP generation from electron transfer, enhanced proton leak, decreased coupling efficiency and redox imbalance associated with chr4p loss. Overall, the metabolic and complex genetic interaction network of chr4p loss may offer valuable insight for treating triple negative breast cancers harboring this chromosome arm deletion and would benefit from the disruption of mitochondria. We also demonstrate the power of systematic identification of genetic interactions using a cohort of cancer cell lines with shared chromosome arm aneuploidies, an approach that can be applied to other chromosome arm aneuploidies and cancer types.

## Methods

### Cell culture

HCC1143, HCC1806, BT-549, HS 578T, MDA-MB-157 and MDA-MB-468 cell lines were cultured in RPMI (Thermo Fisher Scientific, #11875093), 10% fetal bovine serum (FBS) (Thermo Fisher Scientific, #12483020), 50 µg/mL gentamicin (Thermo Fisher Scientific, #15710064). HEK-293T cell line was cultured in DMEM (Thermo Fisher Scientific, #11995073) with 10% FBS. All cell lines used were routinely tested for Mycoplasma (Mycoplasma PCR Detection Kit, abm #G238) and were authenticated using short tandem repeat analysis. All cells were maintained at 37°C, 5% CO_2_.

### Sample classification by chr4p copy number status

#### Primary tumor / patient derived xenografts

Copy number variation analysis of whole genome sequencing (WGS) data for basal breast cancer PT and PDX samples were obtained from a previous study^20^. A sample was classified as chr4p loss if ≥50% of the chromosome arm was deleted, otherwise it was considered chr4p neutral. A receiver-operating characteristic (ROC) analysis was used to identify the relative copy number threshold, represented as log_2_ ratio values that maximized the concordance between PT and PDX chr4p copy number status considering the varying PT sample purity. A true positive referred to PT and PDX samples exhibiting a concordant chr4p copy number status, whereas a false positive referred to PT and PDX samples exhibiting a discordant chr4p copy number status. Log_2_ copy ratio < −0.3 maximized sample concordance and was chosen for all downstream analyses. Samples with ≥ 25% chr4p gain (log_2_ copy ratio > 0.3) were excluded from the analysis.

#### CCLE cell lines

The WGS and whole-exome sequencing (WES) copy number data, RNA sequencing gene expression data and “CRISPR-based gene inactivation data (Achilles gene effect)” of basal breast cancer cell lines were obtained from the DepMap database (https://depmap.org/portal/, 24Q2 release)^31,64,65^. The basal molecular PAM50 subtype of the breast cancer cell lines was manually confirmed by literature curation. A cell line was classified as chr4p loss if ≥50% of the chromosome arm was deleted using log_2_ copy ratio < −0.3, otherwise it was considered chr4p neutral. Cell lines with ≥ 25% chr4p gain (log_2_ ratio > 0.3) were excluded from the analysis. The relative copy number is represented as log_2_ ratio values, which are computed from the log_2_ ratio of the observed signal intensity to the expected signal intensity. Copy ratio genomic heatmaps were generated using the R package ComplexHeatmap^66^.

Long-read sequencing by Oxford Nanopore as described previously of the MDA-MB-157 cell line confirmed chr4p copy neutral status consistent with the analysis of the CCLE cell line copy number data. Briefly, Nanopore long-reads were mapped to the GRCh38 genome using minimap2_v2.17^67^. Clair3_v1.0.8^68^ was used to call single nucleotide variants on a chromosome level (using options: --platform=“ont” and --use_longphase_for_final_output_phasing); the calls were then merged and filtered. Dysgu_v1.6.6^69^ on nanopore mode was used to call large structural variants. QDNAseq_v1.40.0^70^ was used to call copy number variants, where the alignments were binned in 15 kbp long sections.

### Differential gene expression analysis

RNA-seq counts were obtained for CCLE cell lines from DepMap portal (https://depmap.org/portal/, 24Q2 release)^31,64,65^ and for PT/PDXs from a previous study^20^. For all downstream analyses, lowly-expressed genes with an average read count lower than 10 across all samples were excluded. Raw counts were normalized using *edgeR*’s TMM algorithm^71^ and were then transformed to log_2_-counts per million (logCPM) using the *voom* function implemented in the *limma* R package^72^. To assess differences in gene expression levels, a linear model was fitted using *lmfit* function with parameter *method=“robust”*. Nominal p-values were corrected for multiple testing using the Benjamini-Hochberg method. Gene set enrichment analysis based on pre-ranked gene list by t-statistic was performed using the R package *fgsea* (http://bioconductor.org/packages/fgsea/). Default parameters were used.

### Genetic interaction scoring

Pooled genome-wide CRISPR-Cas9 screening data were obtained from DepMap portal (AvanaRawReadcounts.csv; https://depmap.org/portal/, 24Q2 release)^31,65^. MAGeCK Robust Rank Aggreation (RRA) method^32^ was used at default parameters to compare the read counts of the sgRNAs between the chr4p loss (defined by -t) and neutral basal breast cancer cell lines (defined by -c). The MAGeCK software provided output files with the median log_2_ fold change and RRA score per gene. The genetic interactions with a q-value less than 0.1 were retained. Gene set enrichment analysis based on pre-ranked gene list by log_2_ fold-change was performed using the R package *fgsea* (http://bioconductor.org/packages/fgsea/). Default parameters were used.

### STRING data curation

Search Tool for the Retrieval of Interacting Genes/Proteins (STRING) database analysis^40^ was conducted on chr4p negative genetic interaction (MAGeCK score log_2_ fold change <0, *q*<0.1) involved in mitochondria function and redox balance based on STRING database annotations. Interaction networks involving direct physical and indirect functional associations were constructed and visualized by using the String database db version 11. Nodes (proteins) with interaction scores lower than medium confidence level (interaction scores < 0.400) were filtered out and FDR-adjusted p-value threshold was set at 0.05 for all analyses.

### Validation of chr4p negative genetic interactions

For validation of chr4p negative genetic interactions, two sgRNAs per gene with the highest estimated editing efficiency were selected from the Toronto Knockout Library v3 (https://crispr.ccbr.utoronto.ca/crisprdb/public/library/TKOv3/)^73^. Forward and reverse oligos for each sgRNA were designed with overhangs corresponding to the *BsmBI* restriction sites (Supplementary Table 3). sgRNA were cloned into the lentiCRISPRv2 plasmid by Golden Gate Assembly as previously described^74^. Briefly, 25 µL reaction was set up (2.5 µL T7 Ligase Buffer (Cedarlane Labs, #T7DL-100), 1 µL Fast Digest ESP3I (*BsmBI*, Thermo Fisher Scientific # FD0454), 0.25 µL DTT (100 mM, Thermo Fisher Scientific #PR-P1171), 0.125 µL BSA (20 mg/µL, New England Biolabs # B9000S), 0.125 µL T4 ligase (NEB #M0202S), 1 µL 1:10 diluted annealed oligos, 1 µL p70 (25 ng/µL), 19 µL H_2_O) and incubated in a T1000 Thermal cycler (Bio-Rad) for 15 cycles (37°C 5 min, 20°C 5 min). The mix was transformed into NEB stabl *E. coli*, plasmid was purified using miniprep (QIAGEN) and verified by PCR using primers that anneal inside the plasmid backbone with forward primer (5’-3’): AGAATCCTGGAAAG and sgRNA reverse oligo sequence itself for each sample. 25 μL PCR was set up (9.5 μL H_2_O, 1 μL of 25 µM forward primer, 1 μL of 25 µM reverse primer, 1 μL of 25 ng/μL plasmid DNA, and 12.5 μL of DreamTaq DNA Polymerase mix (Thermo Fisher Scientific, #EP0702)). The amplification was initiated with a 2 min denaturation at 95°C, followed by 40 cycles of: 95°C 30 s, 55°C 30 s, 72°C 1 min; the reaction was terminated with a 10 min extension at 72°C. 1.5% agarose gel, 1:10,000 SYBRSafe (Thermo Fisher Scientific, #S33102) was used to visualize the PCR products. Viral particles were produced by co-expressing sgRNA constructs with packaging plasmids psPAX2 (p10) and pMD2.G (p9) in HEK-293T cells using lipofectamine 2000 transfection protocol. Media containing viral particles were collected. Cells were treated with virus in media containing 8 μg/mL polybrene. 24 h after transduction cells were recovered for another 24 h and then HCC1806, MDA-MB-468, MDA-MB-157, BT-549 were selected in 2 μg/mL puromycin dihydrochloride (Sigma, #P7255); HCC1143 in 5 μg/mL and HS 578T in 4 μg/mL for 48 h. MOI of 1 was used for each construct for each cell line. After selection, 20,000 HCC1806, 20,000 HCC1143, 25,000 MDA-MB-468, 40,000 BT-549, 40,000 HS 578T, and 45,000 MDA-MB-157 cells were seeded in 6-well plates for 12 days for the colony formation assay. Cells were fixed with 4% PFA, stained cells with 0.01% crystal violet and quantified manually by comparing to the sg-*AAVS1* control. Confirmed interactions exhibited a greater fitness defect in chr4p loss than neutral cells. Inconclusive interactions refer to inconsistent responses across replicates or between guides.

### High-resolution respirometry assay

A sequential substrate addition protocol was conducted to assess mitochondrial coupled and uncoupled oxygen consumption as well as leak respiration using a two-chamber polarographic sensor (Oxygraph-2k; Oroboros Instruments, Innsbruck, Austria). 10^6^ cells were resuspended in 1 mL Mir05 buffer (Oroboros Instrument, #6010-01) and transferred to the respirometer chambers. MiR05 contains (in mM) 0.5 EGTA, 3.0 MgCl_2_·6H_2_O, 60 K-lactobionate, 20 taurine, 10 KH_2_PO4, 20 HEPES, 110 sucrose, and 1 g/L BSA (pH 7.1). All respiratory measurements were carried out in a hyper-oxygenated environment to avoid oxygen diffusion limitations at 37°C. First, complex I-linked substrates malate (2 mM), pyruvate (6 mM) and glutamate (10 mM) were added to assess leak mitochondrial respiration in the absence of ADP. Subsequently, to measure respiration across complex I in the presence of ADP, a saturating concentration of ADP (5 mM) was added. Succinate (10 mM) was added to assess maximal OXPHOS capacity across complex I and II of the ETS. Oligomycin (2 μg/mL) was added to block complex V. This was followed by FCCP (carbonylcyanide-4 (trifluoromethoxy) phenylhydrazone, 1 μM) to test for uncoupling.

### Western blot analysis

Western blot was conducted to quantify protein abundance of mitochondrial respiratory complexes. Cells were harvested in RIPA lysis buffer (50 mM Tris-HCl, pH 8.0, 150 mM NaCl, 1% Nonidet P-40 (NP-40) substitute, 0.1% SDS, 0.5% sodium deoxycholate, 1 mM PMSF, 1 mM Na_3_VO_4_, 1 mM NaF, 10 μg/mL aprotinin and 10 μg/mL leupeptin, pH 7.4). Lysates were boiled in SDS sample buffer and 50 μg of protein from whole cell lysate were resolved in 4-15% MiniPROTEAN TGX Gel (Bio-RAD, #4561084) using running buffer (20% SDS, Tris, Glycine). Proteins were transferred on Trans-Blot Turbo RTA Mini LF PVDF membranes (Bio-RAD, #1704274) using a Mini Trans-Blot System from Bio-Rad. Total oxidative phosphorylation (OXPHOS) rodent antibody cocktail (1:2000, MitoSciences, #MS604), and Anti-VDAC1/Porin + VDAC3 (1:1000, Abcam, #ab14734) primary antibodies were used. Anti-mouse IgG (NEB, #7076S) and anti-rabbit (NEB, #7074S) secondary antibodies were used at 1:5000 in blocking solution. Immunoreactivity was detected using Clarity Max ECL Substrate (Bio-RAD, #1705062) and visualized using a CDP-STAR chemiluminescence system (Amersham hyperfilm ECL). Total protein abundance per lane was measured using Ponceau S stain (0.1% in 5% acetic acid, Millipore Sigma, #P7170-1L) and used to normalize protein abundance.

### Immunofluorescent staining and analysis

Cells were seeded in 24-well plates on glass coverslip coated with 1 μg/mL fibronectin (Millipore Sigma, #FC010). Following 24 h, cells were fixed with 2% PFA (10 min), permeabilized with 0.1% saponin (20 min) and blocked with 2% BSA (30 min) and then incubated with anti-TOMM20 (1:100, Santa Cruz #sc11415) primary antibody. The primary antibody was visualized with a fluorescent secondary antibody conjugated to Alexa Fluor 488 raised in donkey (1:1000, Thermo Fisher Scientific, #A21206). Nuclei were counterstained with 0.25 ng/mL DAPI (5 min). All steps were performed at room temperature. Images were acquired on the Nikon C2/TIRF confocal laser scanning microscope (Nikon) using a 63X objective.

### Quantification of reactive oxygen species

#### Mitochondrial superoxides - MitoSOX Red

Mitochondrial superoxide levels were measured using MitoSOX Red (Invitrogen, #M36008). Cells were plated at 0.5 × 10^6^ cells per well in 6-well plates and incubated overnight. On the day of the experiment, MitoSOX Red stock solution (5 mM in DMSO) was diluted to 5 µM in pre-warmed HBSS containing calcium and magnesium (HBSS/Ca/Mg). Cells were washed once with pre-warmed HBSS/Ca/Mg and incubated with 1 mL of 5 µM MitoSOX Red per well at 37°C for 20 min. Staining solution was removed, and cells were washed three times with 2 mL prewarmed HBSS/Ca/Mg. Cells were detached with 500 µL prewarmed TrypLE Express for 5 min, and the reaction was quenched with 2 mL prewarmed RPMI containing FBS. Cells were pelleted by centrifugation at 600 × g for 3 min, resuspended in 500 µL prewarmed HBSS/Ca/Mg and analyzed immediately by flow cytometry.

#### General oxidative stress - CM-H_2_DCFDA

General oxidative stress indicator CM-H2DCFDA (Thermo Fisher Scientific, #88593074) was used to measure reactive oxygen species in cells. 15,000 HCC1806, 15,000 MDA-MB-468, 18,000 cells HCC1143, 25,000 BT-549, 35,000 MDA-MB-157 and 45,000 HS 578T cells were seeded in 96-well plate 72 h prior to the experiment day and cultured until 80-90% confluence in RPMI 1640 Medium, no phenol red (Thermo Fisher Scientific, #11835030). Then, cells were treated for 15 min with 5 μM ROS reagent and imaged using the Nikon Eclipse TiE inverted epifluorescence microscope. Images were segmented using Cellpose Live Cell 1 program, flow threshold 0.3, cell dimeter 30 pixels. Fiji (v.2.3., NIH) was used to quantify flourescence.

#### Lipid peroxidation - BODIPY™ 581/591 C11

Lipid peroxidation was assessed by BODIPY 581/591 C11 undecanoic acid (Thermo Fisher Scientific, #11835030). 10^5^ cells were plated in 96-well plate in 200 μL of media with 1 µM BODIPY 581/591 C11 for 30 min at 37°C. The samples were stained with 20 nM DAPI to exclude dead cells and analyzed immediately by flow cytometry (BD FACSMelody).

### Seahorse XF Analysis

Oxygen consumption rate (OCR) and extracellular acidification rate (ECAR) were measured using a Seahorse XFe96 Extracellular Flux Analyzer (Agilent) according to the manufacturer’s protocol. Briefly, 40,000 cells per well were seeded in Seahorse XF96 microplates and allowed to adhere overnight. The culture medium was replaced with Seahorse assay medium (RPMI, Agilent 103576-100) supplemented with 11 mM glucose, 2 mM glutamine and 1 mM pyruvate. Plates were equilibrated in a non-CO_2_ incubator for 1 h prior to the assay. OCR and ECAR were measured at basal conditions and following sequential injections of oligomycin (1.5 µM), FCCP (2 µM), rotenone and antimycin A (0.5 µM each) and 2-deoxyglucose (50 mM). After the assay, cells were fixed with methanol, stained with 300 nM DAPI, and cell numbers were quantified using a Cytation5 imaging system (Biotek) and CellProfiler software. Data were normalized to cell numbers.

---

**Supplementary Table 1**| **PT, PDX, and CCLE gene expression analysis related to figure 2**|. Differential gene expression of chr4p genes in **a**, primary tumours **b**, patient derived xenografts, **c**, CCLE cell lines with deletion or copy-neutral status of chr4p. Transcriptome-wide differential gene expression in **d**, primary tumours **e**, patient derived xenografts, **f**, CCLE cell lines with deletion or copy-neutral status of chr4p. Log_2_ fold change, p-value and adjusted p-value are shown.

**Supplementary Table 2**| **Chr4p loss genetic interaction analysis and validations related to figures 3 and 4. a**, Chr4p genetic interactions obtained from the analysis of DepMap pooled genome-wide CRISPR-Cas9 screens in basal breast cancer cell lines as analyzed by MAGeCK. Log_2_ fold change of RRA score between chr4p loss vs neutral cell lines, p-value and q-value are shown. Negative log_2_ fold change represents negative genetic interactions, and positive log_2_ fold change represents positive genetic interactions. Significant interactions meet threshold of q < 0.1. **b**, Summary of validations results.

**Supplementary Table 3**| **Plasmids list**. Plasmid IDs and descriptions.

**Extended Data Fig 1:**
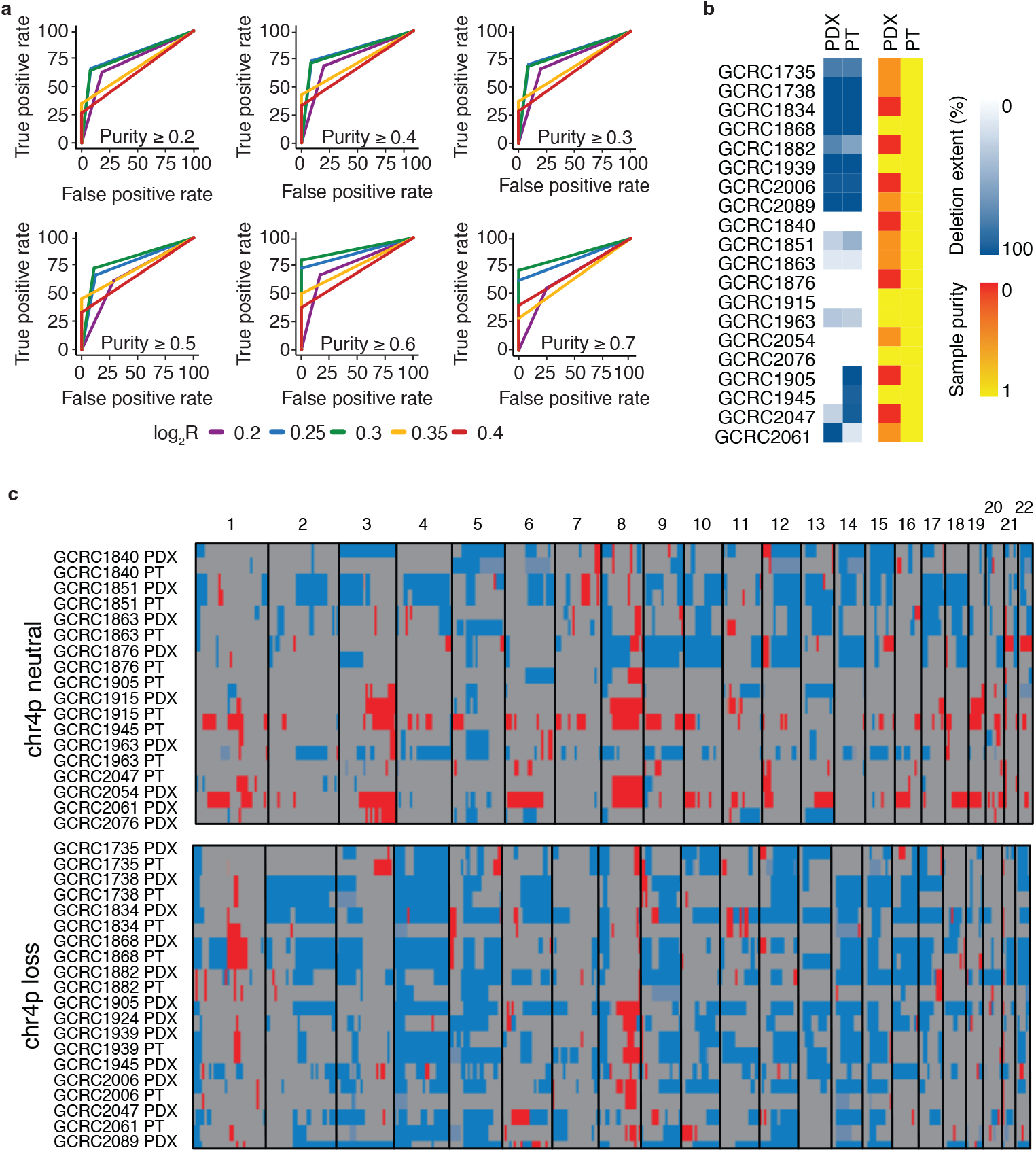
Concordance of copy number profiles between basal breast cancer PT and PDX samples. **a**, A receiver-operating characteristic (ROC) analysis was used to identify the relative copy number threshold, represented as log2 ratio values that maximized the concordance between basal breast cancer primary tumor (PT) and patient-derived xenograft (PDX) chr4p copy number status considering the varying PT sample purity. A true positive referred to PT and PDX samples exhibiting the concordant chr4p copy number status, whereas a false positive referred to PT and PDX samples exhibiting discordant chr4p copy number status. Different colors represent different tested log2 thresholds: purple represents log2 ratio 0.2, blue 0.25, green 0.3, yellow 0.35, red 0.4. **b**, Comparison of deletion extent at log2 ratio 0.3 is shown in white-blue gradient and sample purities in yellow-red gradient for the basal breast cancer PT/PDX samples. **c**, Genome-wide copy number profile for PT/PDX samples, blue represents deletion, grey is neutral state and red is gain. Refer to Methods for details.

**Extended Data Fig 2:**
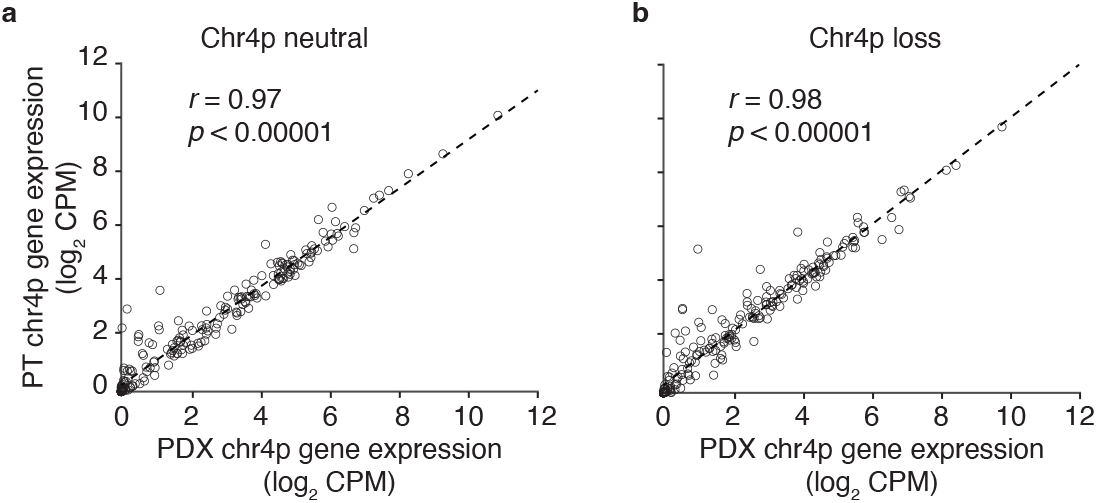
Concordance of gene expression profiles between basal breast cancer PT and PDX samples. **a**, Correlation of chr4p gene expression levels (in log_2_ counts per million (CPM)) between basal breast cancer primary tumor (PT) and patient-derived xenograft (PDX) chr4p neutral and **b, c**hr4p loss samples. The average Pearson correlation coefficient is 0.97-0.98. Refer to Methods for details.

**Extended Data Fig 3:**
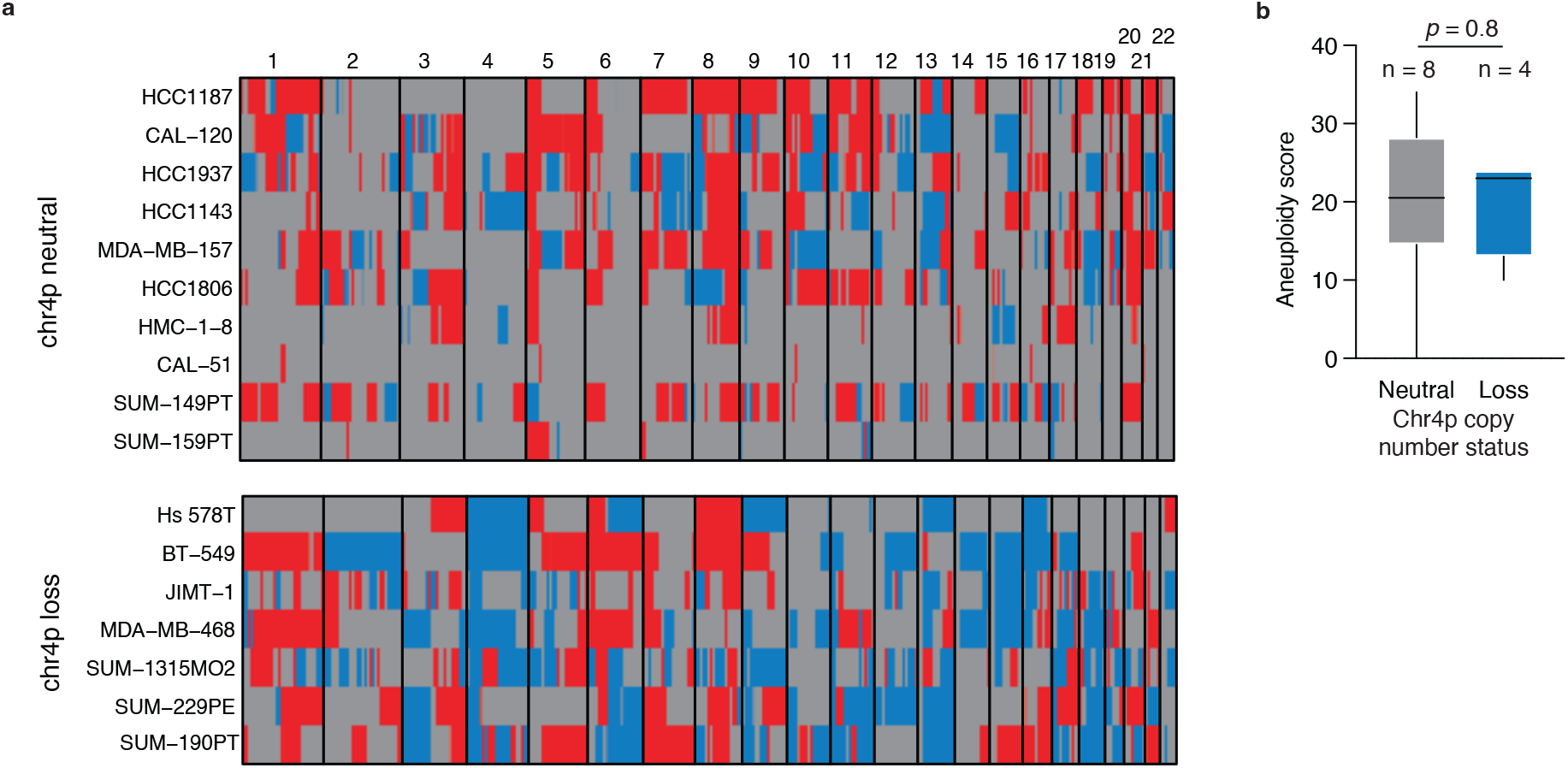
Genome-wide copy number profile for CCLE basal breast cancer cell lines harboring chr4p loss and neutral status. **a**, Genome-wide copy number profile for CCLE basal breast cancer cell lines harboring chr4p loss and neutral status, blue represents deletion, grey is neutral state and red is gain. **b**, Aneuploidy score as reported by Cohen-Sharir et al. shows no statistically significant difference in aneuploidy between chr4p copy neutral vs deletion basal breast cancer cell lines. Significance was assessed by Wilcoxon rank sum test.

**Extended Data Fig 4:**
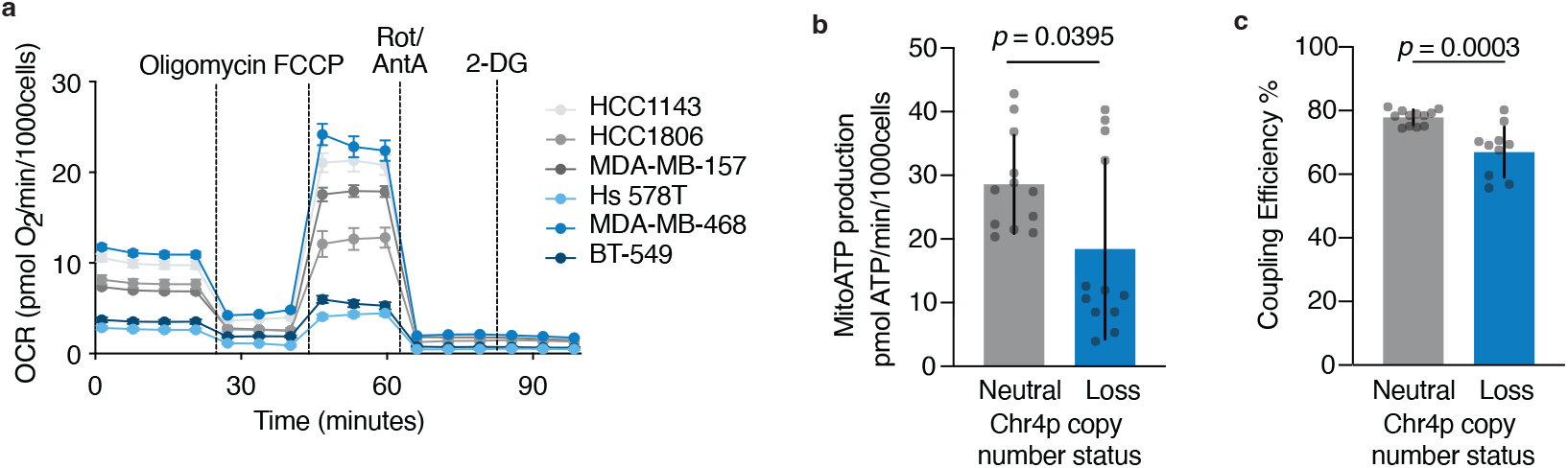
Chr4p loss cells are associated with lower coupling efficiency. **a**, Oxygen consumption rate (OCR) (pmol O2/min/1000 cells). Mean is shown; error bars denote SD. n = 4. **b**, Mitochondrial ATP production was reduced in chr4p loss.Significance was assessed by an unpaired t-test. n=4. Chr4p loss (blue), neutral (grey). **c**, Coupling efficiency was reduced in chr4p loss. Significance was assessed by Wilcoxon rank sum test. n=4. Chr4p loss (blue), neutral (grey).

